# GCN5 restricts PRX-dependent lignin deposition during salt stress in Arabidopsis

**DOI:** 10.64898/2026.01.27.701911

**Authors:** Mansi Sharma, Jan Masood, Pavel Kerchev, Iva Mozgova, Michael Wrzaczek

**Affiliations:** Institute of Plant Molecular Biology, Biology Centre, Czech Academy of Sciences, 37005 České Budějovice, Czech Republic; Faculty of Science, University of South Bohemia, 37005 České Budějovice, Czech Republic; Mendel University in Brno, 613 00 Brno, Czech Republic

**Keywords:** *Arabidopsis thaliana*, *GCN5/HAG1*, histone acetylation, H3K9ac, salt stress, cell wall integrity, lignification, class III peroxidase, reactive oxygen species

## Abstract

Histone acetylation shapes transcriptional programs during environmental stress. The Arabidopsis histone acetyltransferase *GCN5/HAG1*, a catalytic subunit of the SAGA complex, has been implicated in salt stress responses and cell wall integrity. Here, we show that loss of *GCN5* enhances reactive oxygen species (ROS) accumulation under NaCl stress and is accompanied by altered expression of class III peroxidase genes, with strong salt-induced upregulation of *PRX71* and elevated basal *PRX33* transcript abundance. Consistent with a role for these peroxidases in stress-associated cell wall remodeling, overexpression of *PRX71* or *PRX33* in the wild-type is sufficient to promote ectopic lignin deposition in roots. Conversely, *prx71* and *prx33* mutants show improved growth under salt stress and on the cellulose biosynthesis inhibitor isoxaben, and they lack the pronounced ectopic root lignification observed in *gcn5* under salt stress. Chromatin immunoprecipitation followed by qPCR (ChIP-qPCR) reveals reduced H3K9 acetylation at *PRX71* and *PRX33* promoter regions in *gcn5* compared with the wild-type, and reduced transcript abundance of candidate upstream transcription factors (TFs), including *GATA21* and *MYBS2*, accompanied by reduced H3K9ac at their loci. Together, our results support a model in which GCN5 constrains *PRX71/PRX33*-mediated lignification during stress, likely through an indirect regulatory route that integrates chromatin state and transcription factor activity to limit stress-associated lignification while maintaining root growth under salt stress.

## Introduction

Plants are constantly exposed to abiotic and biotic stresses that can occur alone or in combination, requiring rapid signaling as well as longer-term acclimation (Wurzinger, Mair, Pfister & Teige 2011; Boudsocq & Sheen 2013). Among the earliest responses are changes in cytosolic calcium levels and the production of reactive oxygen species (ROS), which act as signaling molecules (Wurzinger *et al*. 2011; Boudsocq & Sheen 2013). Stress also activates mitogen-activated protein kinase (MAPK) and hormone signaling networks (Mittler 2002; Baxter, Mittler & Suzuki 2014; Xia *et al*. 2015; Jalmi & Sinha 2015). Among these stresses, salt stress is a major abiotic constraint because it imposes an early osmotic limitation on growth and a later ionic phase that can exacerbate cellular damage (Munns & Tester 2008). Plants rapidly respond to biotic and abiotic stimuli by triggering responses that involve changes to gene transcription, inherently connected to changes in chromatin structure, and can also orchestrate rapid but also long-lasting epigenetic changes in response to stress. These changes, including histone modifications, are inducible within minutes to hours but can persist for days after the initial stimulus and gradually fade, depending on the locus and stress regime (Ding, Fromm & Avramova 2012; Lämke & Bäurle 2017; Ma, Xing, Li & Jiang 2025). Chromatin structure is determined by combinations of histone modifications including acetylation, methylation, phosphorylation, ubiquitylation, and sumoylation and others, as well as by DNA methylation, which together regulate gene expression without altering the DNA sequence.

Among these modifications, histone acetylation plays a key role in modulating chromatin accessibility. Histone acetylation status is balanced by the activities of histone acetyltransferases (HATs) and histone deacetylases (HDACs), and can be modulated under stress conditions (Ueda & Seki 2020). This balance can be altered, for example, treatment with the HDAC inhibitor Ky-2 increases histone acetylation in *Arabidopsis thaliana* under salt stress, leading to increased expression of salt-responsive genes and contributing to improved salt tolerance (Sako *et al*. 2016). Similarly, MSI1 (WD40-repeat histone binding protein), which forms a complex with HDA19 that deacetylates H3K9 at the abscisic acid (ABA) receptor genes, represses their expression. Loss of MSI1 or HDA19 results in increased ABA sensitivity and enhances tolerance to salt stress in *Arabidopsis* (Mehdi *et al*. 2016). Together, these studies illustrate that modulating histone acetylation plays a key role in controlling stress-responsive gene expression and stress tolerance in plants. One well-characterized HAT module subunit is GCN5 (General Control Non-repressed Protein 5), which is part of the SAGA (Spt–Ada–Gcn5 acetyltransferase) complex, a conserved transcriptional coactivator in eukaryotes that influences a wide range of cellular processes by modulating histones through acetylation and deubiquitination (Moraga & Aquea 2015). In addition to GCN5, SAGA comprises the subunits along with Alteration/Deficiency in Activation 2B (ADA2B), Alteration/Deficiency in Activation 3 (ADA3), SGF29 (SAGA-associated factor 29) subunits (Moraga & Aquea 2015). The catalytic HAT subunit GCN5 acetylates specific lysines on histone H3 at positions K9, K14, K27, and of histone H4 at position K12, and it is crucial for regulating the expression of genes involved in plant development and stress responses (Benhamed, Bertrand, Servet & Zhou 2006; Servet, Conde E Silva & Zhou 2010; Hu *et al*. 2015; Zheng *et al*. 2019; Kim *et al*. 2020). Importantly, GCN5 is involved in salt stress tolerance and cell-wall integrity by modulating H3K9 and H3K14 acetylation linked to the activation of chitinase-like-1 (*CTL1*), polygalacturonase involved in expansion-3 (*PGX3*), and MYB domain protein 54 (*MYB54*) genes associated with cellulose synthesis. This affects turgor, growth dynamics, and cell wall integrity, whereby plants actively remodel cell wall polymers to maintain growth and survival (Zheng *et al*. 2019).

In addition to the SAGA complex, plants have recently been reported to also contain a PAGA (Plant-ADA2A-GCN5-acetyltransferase) complex that influences the transcription of hormone-related genes, controlling plant growth traits such as height and branching (Wu *et al*. 2023). Both SAGA and PAGA share the GCN5 HAT subunit, but the SAGA complex contains ADA2B, while PAGA contains ADA2A. The two complexes independently generate different levels of acetylation, facilitating transcriptional activation (Wu *et al*. 2023).

A key aspect of plant growth and environmental response is the continuous modification of the cell wall, which is loosened by enzymes such as expansins and xyloglucan endotransglucosylase/hydrolase (XTHs) to enable cell expansion. The plant cell wall acts as the plant’s first line of defense against stress during pathogen attack and responds to osmotic or salt stress, and undergoes remodeling to maintain integrity and prevent collapse leading to restricted plant growth (Tenhaken 2015; Cosgrove 2016; Tariq, Ma & Zhao 2025). The plant cell wall consists of cellulose, hemicellulose, and pectin, with cellulose forming long, unbranched chains of β-1,4-linked glucose units (Somerville *et al*. 2004), synthesized by cellulose synthase (CESA) proteins (Cosgrove 2005; Gutierrez, Lindeboom, Paredez, Emons & Ehrhardt 2009). Loss of function or chemical inhibition of CESA by compounds such as by isoxaben or thaxtomin A, disrupts cellulose biosynthesis and cell wall integrity, which results in an ectopic lignin deposition (Caño-Delgado, Penfield, Smith, Catley & Bevan 2003; Persson *et al*. 2007; Bischoff, Cookson, Wu & Scheible 2009). In addition to its ectopic deposition under stress, lignin is normally deposited in the secondary cell wall, strengthening the tissue, making the wall less permeable to water and facilitating water transport through the xylem (Rogers & Campbell 2004; Boyce *et al*. 2004). Abiotic and biotic stresses can modulate the expression of lignin biosynthesis genes. In *Arabidopsis* under salt stress, lignin accumulation is accompanied by higher expression of lignin biosynthesis genes (Moura, Bonine, De Oliveira Fernandes Viana, Dornelas & Mazzafera 2010; Chun *et al*. 2019). The stress-induced lignin deposits occur as a secondary cell wall remodeling response mediated by Class III peroxidases (PRXs), which comprise 73 isoforms in *Arabidopsis* that use H₂O₂ to oxidize monolignols, leading to lignin formation (Passardi, Penel & Dunand 2004; Marjamaa, Kukkola & Fagerstedt 2009; Bischoff *et al*. 2009). Specific PRXs, including *PRX2*, *PRX25*, and *PRX71*, contribute to stem lignification, with mutants showing reduced lignin without developmental defects (Shigeto, Kiyonaga, Fujita, Kondo & Tsutsumi 2013). *PRX71* also plays a role in reinforcing the cell wall, which helps limit cell expansion, and is upregulated in plants with altered pectin or after cellulose synthesis inhibition, promoting ROS accumulation and influencing growth (Raggi *et al*. 2015), while knockdown of *PRX33* and *PRX34* reduces MAMP-induced defense proteins, lowers apoplastic ROS production, and alters cell wall remodeling enzymes (Daudi *et al*. 2012; O’Brien *et al*. 2012). Together, these observations place peroxidase-driven ROS and lignification at the intersection of stress signaling and cell wall remodeling. Yet it remains unclear how the PRX gene expression is tuned at the chromatin-level during salt stress.

Here, we investigate how GCN5 influences the expression of two class III peroxidases, *PRX71* and *PRX33*, and how this relates to ROS accumulation and lignin deposition during salt stress. Using T-DNA insertion mutants and *PRX71* and *PRX33* overexpressing lines, histochemical assays, gene expression analyses, and ChIP-qPCR, we identified an association between GCN5-dependent acetylation and a transcriptional network that constrains peroxidase-linked lignification in stressed seedlings. By connecting a chromatin regulator (GCN5) to distinct PRX loci and downstream cell wall phenotypes, this work extends our understanding of how epigenetic control helps balance stress protection with growth in salt-challenged seedlings.

## Materials and Methods

### Plant material and growth conditions

*Arabidopsis thaliana* T-DNA insertion line *gcn5* (*SALK_048427*) was obtained from the Arabidopsis Biological Resource Centre (ABRC). The T-DNA insertion lines *prx71-1* (*SALK_123643C*), *prx71-2* (*SALK_121202C*), and *prx33* (*SALK_062314C*) were obtained from the Nottingham Arabidopsis Stock Centre (NASC) (Alonso *et al*. 2003). All mutants are in the Columbia-0 (Col-0) background, and Col-0 was used as the wild-type control for all experiments. Primers used in this study are listed in Table S1.

For genotyping, genomic DNA was prepared directly from seedling tissue and PCR was performed using Phire™ Plant Direct PCR Master Mix (Thermo Scientific, F130WH) following the manufacturer’s instructions. Insertion genotypes were confirmed by PCR (Supplementary Figure S4a). For transcript verification of insertion lines, semi-quantitative RT-PCR was performed using total RNA isolated from 10-day-old seedlings, and first-strand cDNA was synthesized as described by (Colina, Krasensky-Wrzaczek, Pérez-Guillén & Wrzaczek 2026). Expression was normalized to the reference gene (*PP2AA3*) (Supplementary Figure S4b).

Seeds were surface sterilized in 70% (v/v) ethanol containing 0.02% (v/v) Triton X-100 for 5 min, followed by 1 min in 96% (v/v) ethanol and five rinses in sterile water. Seeds were plated on half-strength Murashige and Skoog (MS) medium (Duchefa, M0222.0050) adjusted to pH 5.8 and supplemented with 0.8% (w/v) agar and 0.05% (w/v) MES unless stated otherwise. After 2 days of stratification in the dark at 4 °C, plates were transferred to a growth chamber maintained at 22 °C under a 16 h light/8 h dark photoperiod (light intensity 128.26 μmol m-2 s-1). All treatments were performed using sodium chloride (P- Lab, Q09101) and isoxaben (Merck, 36138), unless otherwise indicated.

### Plasmid construction and plant transformation

*PRX71* and *PRX33* overexpression constructs were generated using MultiSite Gateway cloning (Invitrogen; Thermo Fisher Scientific). The coding sequences (CDS) of *PRX71* (984 bp) and *PRX33* (1,062 bp) were amplified from Arabidopsis cDNA. PCR products were recombined into pDONR221 using BP Clonase II enzyme mix (Invitrogen), yielding entry clones carrying *PRX71* or *PRX33* CDS.

For expression in plants, an LR recombination reaction was performed using LR Clonase II Plus enzyme mix (Invitrogen) to assemble the *CaMV 35S* promoter, an mVenus YFP tag, and the *PRX* CDS fragments into the destination vector *pBm43GW* (Karimi, De Meyer & Hilson 2005). The resulting constructs (*35S::PRX71-YFP* and *35S::PRX33-YFP*) were introduced into *Agrobacterium tumefaciens* strain GV3101 (pSoup) (Lifeasible) and transformed into Col-0 plants by floral dip. T1 plants were used for all subsequent experiments.

### Histochemical detection of ROS accumulation

Reactive oxygen species accumulation was assessed using 3,3′-diaminobenzidine (DAB) staining. Ten-day-old seedlings were treated with 0 mM or 150 mM NaCl or 300 nM isoxaben on half-strength liquid MS medium for 6 hours. Seedlings were then infiltrated with 0.1% (w/v) DAB (Sigma-Aldrich, D5637) dissolved in 10 mM MES buffer (pH 6.5) until fully saturated and incubated for 40 min under light. After staining, seedlings were rinsed with water and chlorophyll was removed by incubation in 100% ethanol overnight. Samples were stored in water until imaging. Images were acquired using an Olympus SZ16 stereomicroscope equipped with a PROMICAM 2-5CP digital camera.

### Histochemical detection of lignin deposition

Lignin deposition in roots was assessed by phloroglucinol–HCl staining. Two-weeks-old seedlings were treated with 0 mM or 150 mM NaCl on half-strength liquid MS medium for 6 hours. Roots were stained with 1% (w/v) phloroglucinol–HCl (Sigma-Aldrich, P3502) for 10 min at room temperature. Stained samples were imaged immediately using an Olympus MacroView MVX10 stereomicroscope.

### Root growth assays

For root length measurements, seedlings were grown vertically on half-strength MS medium for 5 days and then transferred to one of the following media: control (0 mM NaCl), 150 mM NaCl, or 2.5 nM isoxaben. Plates were returned to vertical orientation, and seedlings were grown for an additional 7 days. Primary root lengths were measured at the endpoint (day 12). Each treatment was performed using at least three independent biological replicates.

### Survival rate analysis

For salt survival assays, 5-day-old seedlings were transferred to half-strength MS medium containing 0 mM or 150 mM NaCl and grown for 7 days. Survival was determined as the percentage of seedlings that remained green and viable relative to the total number of transferred seedlings. Each treatment included at least three biological replicates.

### RNA extraction and quantitative real-time PCR (qRT–PCR)

For gene expression analyses, seedlings were grown on half-strength MS medium (pH 5.8) without sucrose for 10 days. Salt treatments were performed by transferring seedlings to liquid MS medium containing 150 mM NaCl, and tissue was harvested at 0, 3, 6, and 12 hours after treatment. Samples were immediately frozen in liquid nitrogen and homogenized with glass beads using a bead mill. Total RNA was extracted using ROTIZOL (Karl Roth, 9319.2) according to the manufacturer’s instructions, following the workflow described by (Colina *et al*. 2026). RNA was treated with DNase I to remove genomic DNA contamination as described by (Colina *et al*. 2026). First-strand cDNA was synthesized from 1 µg DNase-treated RNA using the RevertAid cDNA Synthesis Kit (Thermo Fisher Scientific, K1621) with oligo(dT) primers.

Quantitative real-time PCR was performed using gene-specific primers (Supplementary Table S1) with DBdirect PCR SYBR Mix SuperSens (DIANA Biotechnologies, DB-1274). Amplification was measured with a thermocycler (Bio-Rad, CFX Connect Real-Time PCR Detection System). Expression was normalized to the housekeeping gene (*PP2AA3*), following (Czechowski, Stitt, Altmann, Udvardi & Scheible 2005). Relative expression levels were calculated using the ΔΔCT method. Each condition was measured using three independent biological replicates and two technical replicates per biological replicate.

### Chromatin immunoprecipitation (ChIP) and ChIP–qPCR

ChIP was performed using a crosslinked Arabidopsis ChIP protocol adapted from (Mozgová *et al*. 2015). Wild-type (Col-0) and gcn5 seedlings were grown on half-strength MS plates for 10 days and treated in liquid half-strength MS with 150 mM NaCl for 3 hours; control samples were mock-treated without NaCl. Tissue (∼100 mg) was crosslinked in 1% formaldehyde by vacuum infiltration (10 min) and quenched with 0.125 M glycine (5 min). Samples were washed, snap-frozen, and ground in liquid nitrogen. Nuclei were extracted in Modified Extraction Buffer, filtered through Miracloth, and pelleted (1500 rcf, 10 min, 4 °C). Nuclei were lysed in 1% SDS and chromatin was fragmented by sonication (Diagenode Bioruptor 300, 7 cycles, 30 s ON/30 s OFF, high power) to obtain ∼0.3–1.5 kb fragments, then diluted with ChIP dilution buffer to reduce SDS to 0.1% and cleared by centrifugation. Ten percent of each sample was saved as input DNA. Immunoprecipitation was performed overnight at 4 °C with rotation using anti-H3K9ac (Sigma-Aldrich, 07-352) and, where indicated, total H3 (Sigma-Aldrich, 07-690) and IgG (Sigma-Aldrich, R2655) as a negative control. Immune complexes were captured using Protein A magnetic beads and washed using low- and high-salt wash buffers. Chromatin was eluted twice (elution buffer; 65 °C, 15 min) and crosslinks were reversed by NaCl addition and incubation at 65 °C for ≥6 hours. Samples were treated with proteinase K (45 °C, 1 hour), and DNA was purified using the IPure kit v2 (Diagenode, C03010015).

ChIP qPCR was done using DBdirect PCR SYBR Mix SuperSens (DIANA Biotechnologies, DB-1274), and measured with a thermocycler (Bio-Rad, CFX Connect Real-Time PCR Detection System). ChIP enrichment was quantified by qPCR using three biological replicates, with two technical replicates per biological replicate. Enrichment was calculated relative to input DNA and normalized to total H3. A transposable element locus (*At4g03770*) was used as an H3K9ac negative control locus.

### Statistical analysis

All statistical analyses were performed in R (2023-12-1; Build 402; Posit Software) using RStudio (2024-04-24, 4.4.0; R Foundation). Data were initially tested for normality and homogeneity of variances. Levene’s test and F-test were applied to assess variance homogeneity. For datasets meeting parametric assumptions, *t*-tests, and two-way ANOVA were used, with Tukey’s post-hoc test applied for multiple pairwise comparisons. For datasets that did not meet parametric assumptions, Kruskal–Wallis tests were used as a non-parametric alternative followed by wilcoxon post-hoc test for pairwise comparisons. In all analyses, *p* < 0.05 was considered significant.

## Results

### Loss of GCN5 increases ROS accumulation and alters *PRX33* and *PRX71* expression during salt stress

To confirm and extend the previous reports that *gcn5* is sensitive to salt stress (Zheng *et al*. 2019), we analyzed the survival rate and root growth of *gcn5* seedlings under 150 mM NaCl. We included *gcn5* mutants, wild-type plants, and three independent complementation lines expressing *pGCN5::GCN5-GFP* in *gcn5* mutant background (described in De Smet *et al*. 2025) to confirm that the observed phenotypes were specifically caused by the loss of GCN5. Survival rates and primary root lengths of plants grown on 150 mM NaCl were measured on day 12 and normalized to control conditions (Figure S1a, b). The *gcn5* mutant showed reduced survival (Figure S1a) and shorter primary root length (survival rate, Figure S1a; root length, (Figure S1b) compared with wild-type. All three complementation lines partially rescued the survival phenotype (Figure S1a) and restored root length to levels similar to the wild type (Figure S1b). Thus, consistent with previous findings (Zheng *et al*. 2019), we observed that the *gcn5* mutant analyzed in this study was sensitive to salt stress.

Salt stress induces ROS accumulation (Mittler 2002) and GCN5 may play a role in maintaining ROS balance under salt stress. TaHAG1, a wheat ortholog of GCN5, contributes to ROS balance under salt stress (Zheng *et al*. 2021). We next investigated whether the loss of GCN5 affects ROS accumulation in *Arabidopsis.* To examine this, we performed DAB staining to visualize hydrogen peroxide (H_2_O_2_) accumulation in 10-day-old *gcn5* and wild-type plants following 6 hours of treatment with 150 mM NaCl or 300 nM isoxaben, both of which can trigger ROS accumulation. Isoxaben triggers a wide spectrum of cell wall integrity-related adaptations, including hormone signaling and lignin accumulation, by disrupting cellulose biosynthesis (Scheible, Eshed, Richmond, Delmer & Somerville 2001; Denness *et al*. 2011; Hamann 2015). Under mock conditions, ROS accumulation was not detectable in either wild-type plants or the *gcn5* mutant. However, in response to NaCl and isoxaben treatments, ROS accumulation was enhanced in the *gcn5* mutant compared with wild-type (Figure 1a).

**Figure 1.**
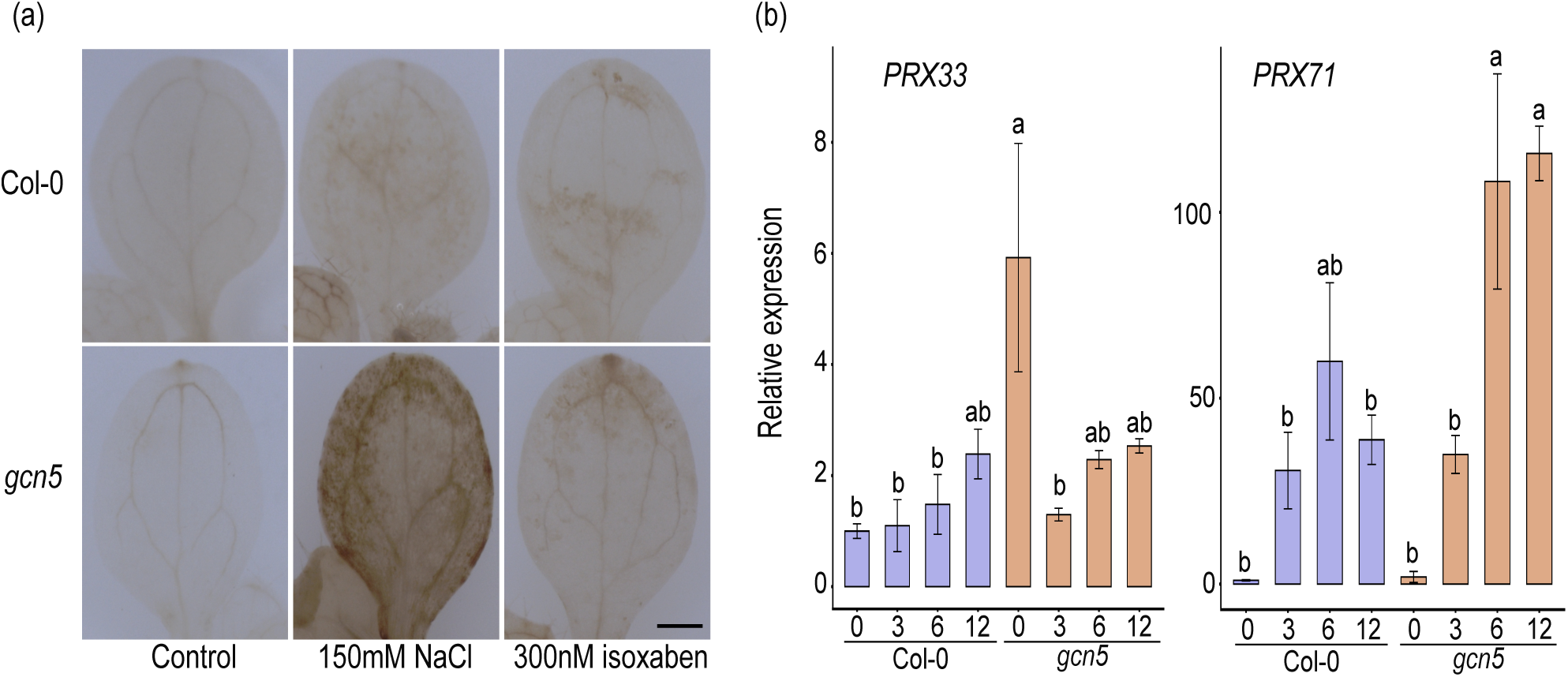
Enhanced ROS accumulation and upregulation of *PRX* genes in the *gcn5* mutant. (a) DAB staining of 10-day-old wild-type and *gcn5* seedlings after 6 hours of treatment under control conditions, 150 mM NaCl, or with 300 nM isoxaben. Images are representative of at least three independent biological experiments. Scale bar = 0.5 mm. (b) Expression levels of selected PRX genes at different time points during 150 mM NaCl treatment. Transcript levels were normalized to a housekeeping gene and expressed relative to wild type at 0 hour (=1). Data represent mean ± SE of three biological replicates. Statistical significance was determined using two-way ANOVA followed by Tukey’s HSD test. Different letters indicate significant differences.

To explore the molecular basis of the elevated ROS accumulation in *gcn5* under stress conditions, we analyzed publicly available transcriptomic data to identify genes differentially expressed in the *gcn5* mutant (Wu *et al*. 2023). In this dataset, 23 peroxidase genes were misregulated in the *gcn5* mutant, of which 20 were upregulated and 3 downregulated (Figure S1c). Since the enhanced salt sensitivity of the *gcn5* plants was previously connected with altered cell wall synthesis and enhanced lignin deposition (Zheng *et al*. 2019), we focused on class III PRXs involved in lignin polymerization (Cosio & Dunand 2009). The upregulation of class III PRX transcripts in *gcn5* could influence ROS balance, as they utilize H_2_O_2_ during lignin polymerization. We hypothesized that the increased expression of these PRX genes in the *gcn5* mutant contributes to lignin deposition, as lignin is known to accumulate in response to cell wall damage (Passardi *et al*. 2004; Denness *et al*. 2011). Based on this, *PRX2*, *PRX25*, *PRX71*, and *PRX52* were selected for their reported roles in lignification (Shigeto *et al*. 2013; Fernández-Pérez, Pomar, Pedreño & Novo-Uzal 2015; Shigeto & Tsutsumi 2016). In addition, *PRX33* was included based on its reported involvement in ROS-related responses (Daudi *et al*. 2012; Arnaud *et al*. 2017), in line with the observed ROS accumulation in the *gcn5* mutant.

Transcript abundance of these selected PRX genes were quantified by qRT-PCR at different time points after treatment with 150 mM NaCl in wild type and *gcn5* (Figure S1d). Among them, *PRX71* and *PRX33* showed consistently higher expression in *gcn5* compared to wild-type plants. *PRX33* was elevated in *gcn5* under control conditions but showed a weaker induction upon salt treatment compared to wild type. By contrast, *PRX71* transcript abundance increased after 6 hours of NaCl treatment and continued to rise up to 12 hours, reaching levels significantly higher than those in wild type (Figure 1b). Overall, this indicates that *PRX33* and *PRX71* are upregulated in the *gcn5* mutant and were therefore selected for subsequent functional analyses.

### *PRX71* and *PRX33* overexpression alters lignin deposition

To test whether the upregulation of these PRX genes is sufficient to promote lignin deposition, we next examined *PRX71* and *PRX33* overexpression lines together with the *prx33*, *prx71*, and *gcn5* mutant lines. Based on our observation that the loss of GCN5 resulted in enhanced *PRX71* and *PRX33* expression and the known involvement of class III PRXs in lignin polymerization, we next investigated whether upregulation of *PRX71* and *PRX33* contributes to enhanced lignin deposition and potentially to maintaining cell wall integrity. To visualize lignin, we used phloroglucinol-HCl staining, which reacts with the aldehyde group of lignin and results in a pink coloration (Davidson *et al*. 1995; Pradhan Mitra & Loqué 2014). The *gcn5* mutant displayed increased lignin deposition in the vasculature compared with wild-type roots, whereas *prx71* and *prx33* resembled wild-type roots. *PRX71* and *PRX33* overexpression lines showed enhanced lignin deposition in the vasculature, similar to *gcn5,* but in addition displayed ectopic lignin accumulation (Figure 2). These results indicate that elevated PRX expression is sufficient to promote vascular and ectopic lignin deposition, recapitulating key aspects of the *gcn5* phenotype, independent of GCN5 loss.

**Figure 2.**
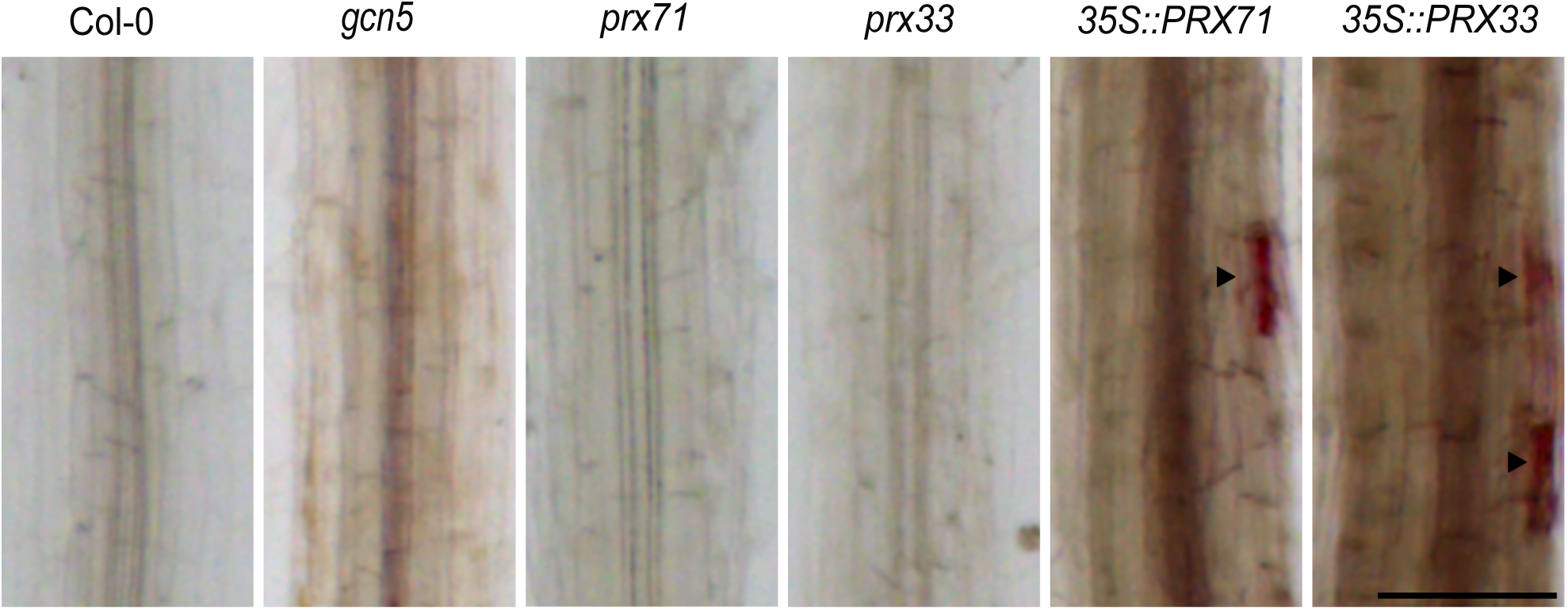
*PRX71* and *PRX33* contribute to lignin deposition. Lignin deposition in roots of wild type, *gcn5* mutant, *prx71* mutant, *prx33* mutant, and *35S::PRX33* and *35S::PRX71* overexpression lines under control conditions. Arrows indicate ectopic lignin deposition. Images are representative of at least three independent biological experiments. Scale bar = 50 μm.

Given these observations and because GCN5 loss leads to reduced cellulose content, as it regulates cellulose synthesis genes (Zheng *et al*. 2019), increased lignin deposition could serve as a compensatory reinforcement mechanism to maintain cell wall strength. Consistent with this idea, our findings indicate that *PRX71* and *PRX33*, also upregulated in *gcn5*, promote vascular and ectopic lignin deposition, which may help to counteract cell wall disruption.

### Altered lignin deposition and root length responses under salt stress

Considering the increased lignin deposition in both *gcn5* and PRX overexpression lines, we next asked how these changes relate to salt-driven root growth phenotypes. To understand the roles of *PRX71* and *PRX33* in response to salt stress, we analyzed salt sensitivity and root growth following salt stress or isoxaben treatment in the *prx71-1*, *prx71-2*, and *prx33* loss-of-function mutants, together with *gcn5* and wild-type plants. As observed previously (Figure S1a), under salt stress, the *gcn5* mutant showed reduced survival compared to wild-type plants, whereas the complementation lines showed higher survival than the *gcn5*, partially rescuing the phenotype (Figure 3a, S2a). In contrast, the *prx71* and *prx33* mutants showed statistically higher survival than wild-type plants, and *prx71-2* showed a survival rate comparable to *prx71-1* and *prx33* (Figure 3a, S2a). To assess growth responses under salt and cell wall stress, we analyzed the effect of treatment on root length under NaCl and isoxaben treatments (Figure 3b, c). Root lengths were normalized to control conditions in the respective genotype (absolute root lengths are shown in Figure S2b, c). The treatment had a milder effect on root length in the *prx* mutants compared to wild-type. The *gcn5* mutant displayed higher sensitivity, which was restored by GCN5 complementation (Figure 3b, c).

**Figure 3.**
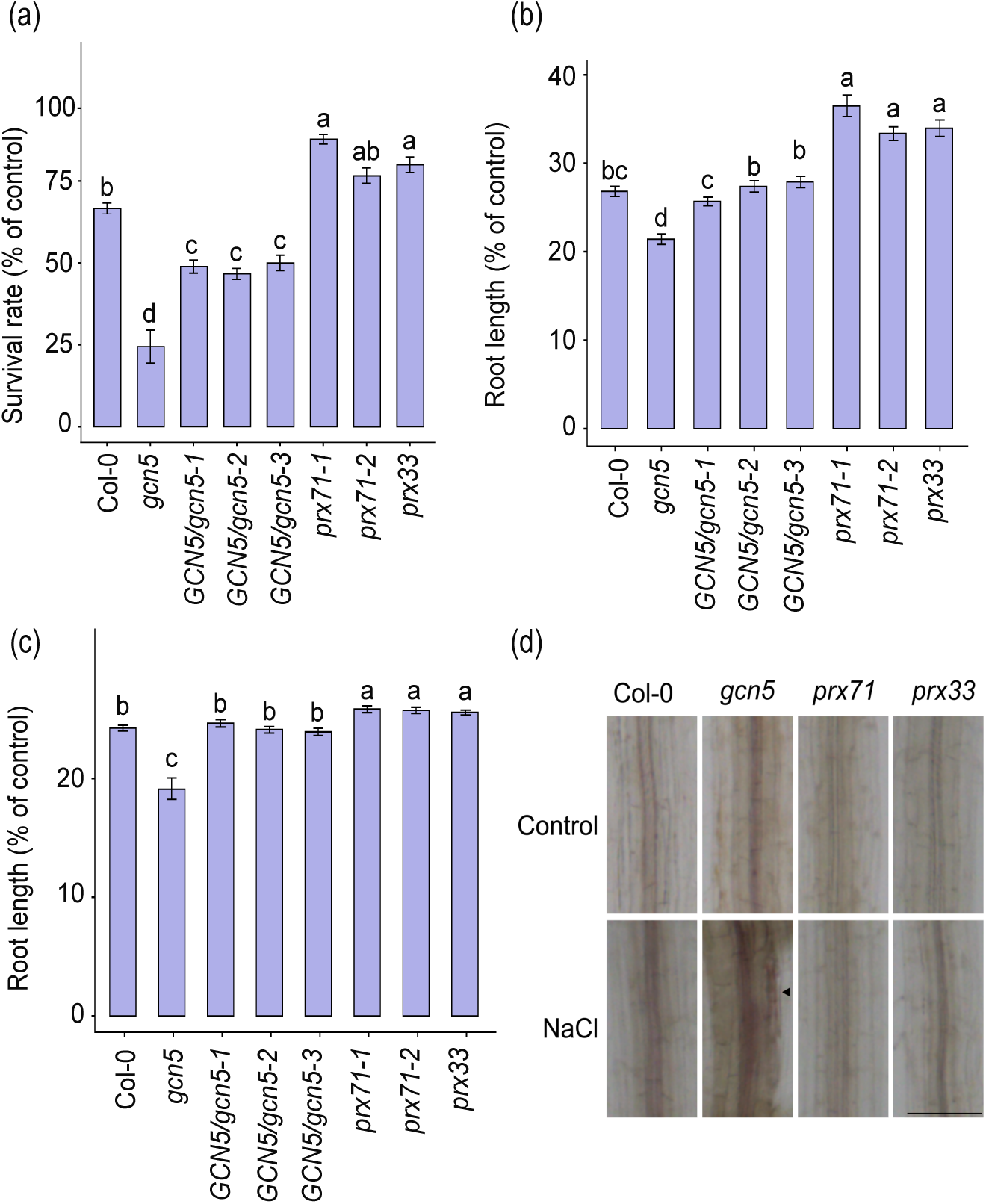
Salt stress responses and lignin deposition. (a) Survival rates under salt stress. Survival was calculated relative to plants grown under control conditions. Data represent mean ± SE (n ≥ 3 biological replicates). Statistical analysis was performed using two-way ANOVA after confirming homogeneity of variance (Levene’s test, p = 0.070), followed by Tukey’s HSD test. Different letters indicate significant differences. (b) Root length was calculated relative to plants grown under control conditions. Primary root length (%) measured at day 12 for seedlings grown on control or 150 mM NaCl containing medium. (c) Primary root length (%) measured at day 12 for seedlings grown on control or 2.5 nM isoxaben containing medium. Root length was calculated relative to plants grown under control conditions. Data in (b) and (c) were analyzed using the Kruskal–Wallis test due to unequal variances (Levene’s test, *p* < 0.05), followed by the Wilcoxon post-hoc test. Different letters indicate significant differences. Absolute root lengths are shown in Figure S2b and S2c. (d) Lignin deposition in roots of wild type, *gcn5*, *prx71*, and *prx33* mutants after 6 hours of 150 mM NaCl treatment. Arrows indicate ectopic lignin deposition in the *gcn5* mutant. Images are representative of at least three independent experiments. Scale bar = 50 μm.

We next examined lignin deposition in 10-day-old roots of *gcn5*, *prx71-1*, and *prx33* mutants in response to salt stress. The *gcn5* mutant showed increased lignin deposition after 6 hours of treatment with 150 mM NaCl, including enhanced accumulation in the vasculature and ectopic deposition. In the wild-type roots, lignin accumulation was also observed after 6 hours of treatment, whereas *prx71-1* and *prx33* mutants did not show ectopic lignin deposition. The absence of lignin deposition in *prx71-1* and *prx33* under salt stress suggests that both these peroxidases are critical for lignin deposition during salt-induced cell wall stress (Figure 3d). Together, these results indicate that *PRX71* and *PRX33* contribute to lignin deposition in response to salt stress. Furthermore, the absence of lignin deposition in *prx71-1* and *prx33* mutants correlates with reduced impact of salt treatment on root length, suggesting that lignin accumulation contributes to root growth restriction in response to stress-associated cell wall disruption.

### Indirect regulation of *PRX71* and *PRX33* transcription by GCN5 through chromatin modification

Given this connection between GCN5, PRX function, and lignin deposition, we next examined whether GCN5 influences *PRX71* and *PRX33* transcription through chromatin modifications. Since *PRX71* and *PRX33* were upregulated in the *gcn5* mutant (Figure 1b), we examined whether this may be reflected in the amount of histone acetylation at the gene loci, in particular at the promoter regions. To this end, we performed chromatin immunoprecipitation (ChIP) assays using an anti-H3K9ac antibody, analyzing histone acetylation at the promoter regions *PRX71* and *PRX33*. Ten-day-old wild-type and *gcn5* seedlings were treated with 150 mM NaCl for 3 hours, at which point we previously observed changes in *PRX* expression (Figure 1b). In line with the histone acetyltransferase function of GCN5, we found reduced H3K9ac enrichment at the promoter regions of *PRX71* and *PRX33* in the *gcn5* mutant compared with wild-type under both control and salt-stress conditions (Figure 4a). In addition, the increase in H3K9ac at *PRX71* upon salt treatment in wild-type was not reflected in *gcn5* (Figure 4a). The increase of H3K9ac at *PRX71* in wild type corresponded with its transcriptional activation upon salt treatment (Figure 1b). However, the expression of *PRX33* and *PRX71* was higher under control (*PRX33*) or salt-treated (*PRX71*) conditions in *gcn5* (Figure 1b) despite the decrease in H3K9ac at both loci in *gcn5*. This suggested that the upregulation of these genes in *gcn5* may be an indirect effect of GCN5 absence. Therefore, we set out to identify potential transcriptional repressors of the PRX genes, hypothesizing that the absence of GCN5 and resulting lower level of H3K9ac at a repressor gene may promote (prime) the transcription of *PRX33* and *PRX71*. Using publicly available transcriptomic and ChIP-seq data, we searched for genes downregulated in *gcn5* that carry less H3K9ac in *gcn5* (Benhamed *et al*. 2008; Wu *et al*. 2023) and that have been identified as direct targets of GCN5 in the wild-type (Kim *et al*. 2020; Wu *et al*. 2023) (Figure S3a). Next, we analyzed potential transcription factor binding sites in the promoters of *PRX71* and/or *PRX33* using the Plant Promoter Analysis Navigator (PlantPAN) 4.0 (Chow *et al*. 2024). This analysis identified transcription factors predicted to bind to one or both genes (Table S2). We then compared these transcription factors, predicted to bind the promoters of *PRX71* and *PRX33*, with the list of GCN5-dependent genes identified from transcriptomic and ChIP-seq data (Figure S3b).

**Figure 4.**
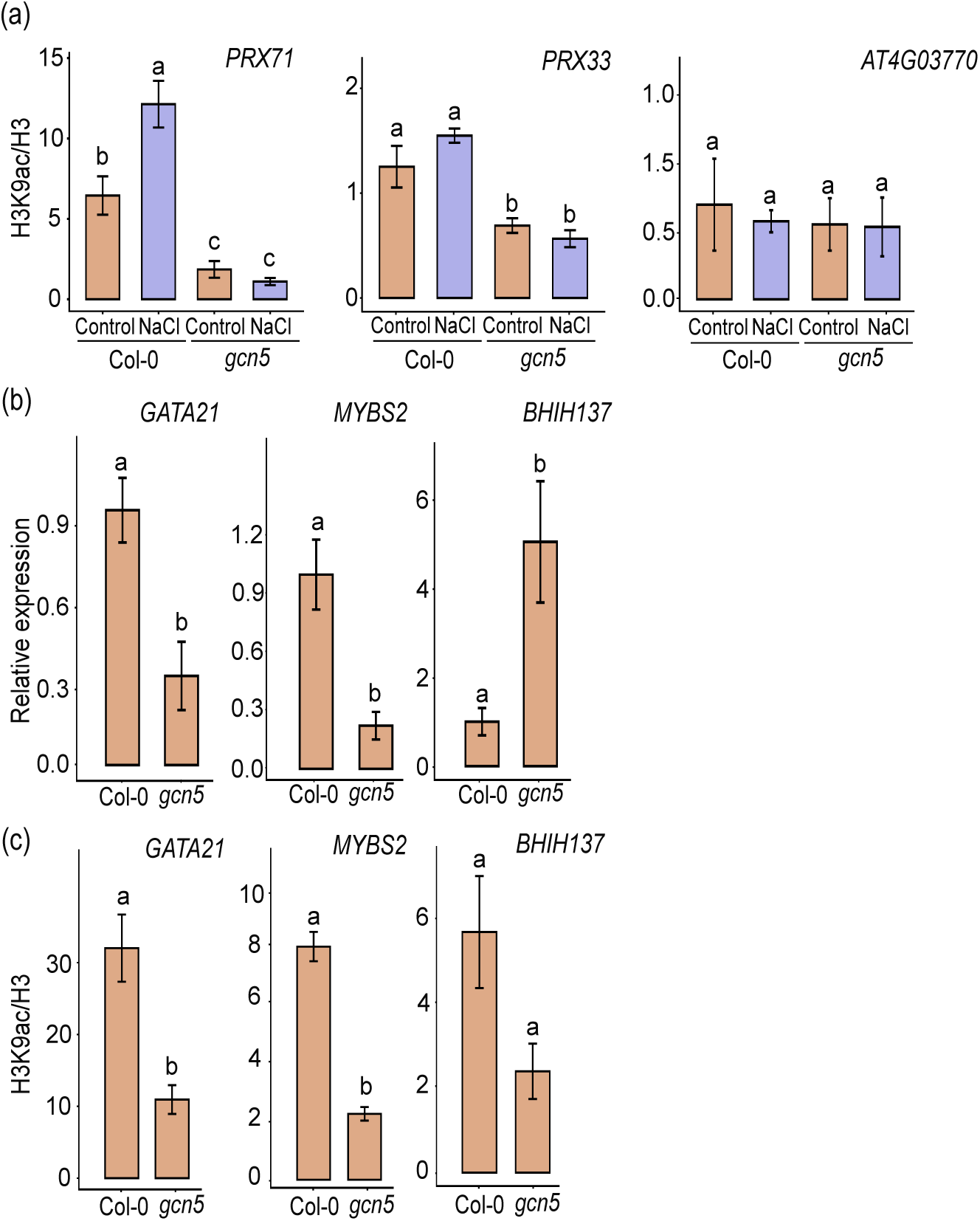
GCN5 indirectly regulates *PRX33* and *PRX71* through chromatin modification and transcription factor activity. (a) ChIP–qPCR analysis of H3K9ac enrichment at the promoter regions of *PRX71* and *PRX33* in wild-type and *gcn5* seedlings under control and 150 mM NaCl conditions. ChIP was performed using anti-H3K9ac antibodies, and enrichment was normalized to total H3. A transposable element locus (*At4g03770*) was included as a negative control. Statistical analysis was performed using two-way ANOVA after confirming homogeneity of variance (Levene’s test, *p* > 0.05), followed by Tukey’s HSD test. (b) Relative expression of candidate transcription factor genes in wild-type and *gcn5* seedlings under control conditions, measured by qRT–PCR. Statistical analysis was performed using a two-tailed Student’s t-test. (c) ChIP–qPCR analysis of H3K9ac enrichment at the promoter regions of *GATA21* and *MYBS2* in wild-type and *gcn5* seedlings under control conditions. ChIP was performed using anti-H3K9ac antibodies, and enrichment was normalized to total H3. Statistical analysis was performed using a two-tailed Student’s t-test.

This led to the identification of transcription factors that could potentially regulate (repress) the *PRX71* and *PRX33* genes in a GCN5*-*dependent manner, including 16 TFs that were either common to both promoters or specific to either *PRX71* or *PRX33* (Figure S3b, Table S2). We next analyzed the transcript levels of the 16 TFs in wild-type and *gcn5* seedlings under control conditions by qRT–PCR (Figure S3c) and identified three candidates that were differentially expressed in *gcn5* (Figure 4b, S3c). Among these, GATA-type zinc finger transcription factor 21 (*GATA21*) was predicted to bind the promoters of both *PRX71* and *PRX33*, whereas MYB-related transcription factor 2 (*MYBS2*) and basic helix-loop-helix 137 *(BHLH137*) were predicted to bind the *PRX33* promoter. *GATA21* and *MYBS2*, but not *BHLH137*, were downregulated in *gcn5*, in line with the requirement of GCN5 for their activation (Figure 4b). We next asked whether the downregulation of these genes in *gcn5* is associated with a loss of H3K9ac. Accordingly, we observed lower enrichment of H3K9ac at the promoters of *GATA21* and *MYBS2* (Figure 4c). Together, these findings suggest that GCN5 may influence *PRX71* and *PRX33* gene expression indirectly through transcription factors such as *GATA21* and *MYBS2*.

## Discussion

GCN5 is a subunit of the histone acetyltransferase (HAT) module present in the SAGA and PAGA complexes, where it facilitates histone acetylation, thereby contributing to the regulation of gene expression and numerous plant developmental pathways (Vlachonasios, Thomashow & Triezenberg 2003; Wu *et al*. 2023). Consistent with this central regulatory role, loss of GCN5 function results in smaller rosettes (Vlachonasios *et al*. 2003), short roots and defects in meristem activity (Kornet & Scheres 2009; Servet *et al*. 2010) as well as other developmental defects, and it is associated with altered expression of stress-responsive genes under various environmental stimuli, such as the heat-induced *HSFA3* and *UVH6* (Benhamed *et al*. 2006; Hu *et al*. 2015). Taken together with our results, these studies position GCN5 as a key coordinator of growth and stress adaptation, and our data extend this role to ROS homeostasis and PRX-dependent lignification during salt stress.

Notably, recent biochemical and genomic analyses have established that GCN5 operates not only within the conserved SAGA complex (containing ADA2B) but also in a plant-specific complex, named PAGA, which incorporates ADA2A and four additional plant-specific subunits (Wu *et al*. 2023). SAGA and PAGA independently generate high and moderate levels of H3K9 acetylation, respectively, and can antagonistically regulate transcription at shared targets (Wu *et al*. 2023). The *gcn5* mutation eliminates the catalytic subunit shared by both complexes.

Thus, the observed reduction in H3K9ac at both PRX and TFs could reflect a combined loss of SAGA and PAGA activity. However, analysis of publicly available ChIP-seq datasets (Wu *et al*. 2023) indicates that *PRX71* and *MYBS2* exhibit reduced H3K9ac in *gcn5* and are direct targets bound by the SAGA component ADA2B, supporting a role in this regulatory pathways (Table S3). Nevertheless, disentangling the relative contributions of SAGA and PAGA to PRX regulation, for example, by comparing *ada2a* and *ada2b* single mutants — could reveal whether the indirect regulatory circuit identified here operates primarily through one or both GCN5-containing complexes. Such analysis could also clarify whether the moderate acetylation typically deposited by PAGA is sufficient to maintain expression of the PRX-constraining TFs *GATA21* and *MYBS2* in the absence of SAGA, or whether SAGA-level acetylation is required.

Previous research showed that in response to salt stress, TaHAG1, the wheat ortholog of *Arabidopsis* GCN5, helps to maintain ROS balance. In TaHAG1 overexpression lines, cytosolic enzymatic antioxidant activity was increased. However, whether this results from direct gene regulation by TaHAG1 or from its effect on ROS-detoxifying pathways remains unclear (Zheng *et al*. 2021). In hexaploid wheat, TaHAG1 directly induces the expression of respiratory burst oxidase (*Rboh*) genes by promoting H3K9 and H3K14 acetylation at their promoters, resulting in increased NADPH oxidase-dependent ROS production that contributes to Na^+^ exclusion and salt tolerance (Zheng *et al*. 2021). By contrast, we observed that loss of Arabidopsis GCN5 led to elevated apoplastic ROS (Figure 1a), driven by class III peroxidases rather than NADPH oxidases. Notably, these findings may reflect compartmentalized ROS management: RBOH-generated superoxide at the plasma membrane might serve as a signaling molecule promoting ion homeostasis, whereas class III PRX-generated H_2_O_2_ in the apoplast might drive monolignol oxidation and lignin polymerization. In addition, GCN5 contributes to maintaining cell wall integrity under salt stress in Arabidopsis by influencing the expression of genes involved in cellulose synthesis (Zheng *et al*. 2019). Compared with wild-type seedlings, *gcn5* exhibits reduced cellulose content, which is further decreased under salt stress, while lignin accumulation is enhanced together with ectopic lignification (Zheng *et al*. 2019). Our work shows that in response to stress, *gcn5* seedlings accumulated elevated levels of ROS and interestingly, showed altered expression of several class III PRXs. Specifically, we identified *PRX71* and *PRX33* as promising candidates (Figure 1b). Previous studies have shown that in *Arabidopsis*, class III PRXs like *PRX71* limit root elongation (Raggi *et al*. 2015) and overexpression of *AtPRX37* leads to a dwarf phenotype (Pedreira, Herrera, Zarra & Revilla 2011), suggesting that altered peroxidase activity may contribute to root growth restriction in *gcn5*. Accordingly, the elevated ROS levels (Figure 1a) and altered *PRX* regulation (Figure 1b, 4a) in *gcn5* suggest that a chromatin-level perturbation is translated into a cell wall-centered growth phenotype under salt stress.

Lignin deposition is a common response to stress-triggered cell wall damage and is considered to enhance cell wall integrity (Caño-Delgado *et al*. 2003; Denness *et al*. 2011). Class III PRXs are centrally involved in controlling lignin deposition (Cosio & Dunand 2009; Marjamaa *et al*. 2009). Enhanced lignin deposition in response to salt stress has been reported in plant species, such as maize (Oliveira *et al*. 2020) and in *Arabidopsis,* showing elevated lignin accumulation (Chun *et al*. 2019). Both *35S::PRX71* and *35S::PRX33* lines exhibited enhanced lignin accumulation similar to the *gcn5* mutant under control conditions. Overall, our data support a role for *PRX71* and *PRX33* in mediating lignification as part of the cell wall reinforcement response. Loss of GCN5 leads to a balance shift towards ROS accumulation and *PRX*-mediated lignification (Figure 1a; Figure 2; Figure 3d). This enhanced lignin deposition correlates with the reduced root growth in response to salt stress (Figure 3b, c). Alternatively, the upregulation of *PRX* expression might be secondary to ROS or cell wall integrity signaling. However, the recapitulation of the enhanced lignification in plants overexpressing *PRX71/PRX33* (Figure 2) suggests that increased PRX abundance and activity are sufficient to drive lignin deposition under non-stress conditions (Figure 2). Together, our results support a functional link between elevated *PRX71*/*PRX33* activity and lignin deposition. Notably, *prx71* and *prx33* loss-of-function mutants showed improved survival under salt stress compared to wild type (Figure 3a), suggesting that elevated *PRX*-mediated lignification can be detrimental to salt tolerance. Consistent with this, *prx71* and *prx33* mutants lacked ectopic lignin deposition under salt stress (Figure 3d), supporting a model in which restricting stress-induced lignification is associated with improved survival. Practically, these results suggest that modulating *PRX* activity, or upstream regulators such as GCN5-dependent pathways, may offer a route to improve salt stress resilience in crops.

Loss of GCN5 simultaneously reduces cellulose synthesis and derepresses *PRX71/PRX33* expression, promoting ectopic lignin deposition (Zheng *et al*. 2019). Together, these parallel effects point to GCN5 as a chromatin-level switch involved in coordinating the balance between primary (cellulose-based) and secondary (lignin-based) cell wall composition. In wild-type plants under salt stress, GCN5-dependent acetylation likely maintains cellulose production to preserve primary wall integrity while simultaneously limiting compensatory lignification. This balance is disrupted in *gcn5* and the reduced cellulose content could trigger cell wall integrity sensing pathways that amplify *PRX*-mediated lignin deposition, resulting in cell wall stiffening at the cost of growth. In the future, simultaneously restoring cellulose synthesis and suppressing lignification could clarify whether these two arms of the *gcn5* cell wall phenotype are additive or epistatic.

While GCN5 is widely recognized for its role in histone acetylation, recent work in *Arabidopsis* has shown that it can also acetylate the transcriptional co-repressor TOPLESS (TPL) and thereby contribute to gene repression. This acetylation promotes interaction of TPL with NOVEL INTERACTOR OF JAZ (NINJA) and its recruitment to target promoters, resulting in chromatin compaction and transcriptional repression (An *et al*. 2022). In our study, reduced H3K9ac enrichment at the *PRX71* and *PRX33* promoter regions in the *gcn5* mutant argues against a simple model in which GCN5 directly activates these PRX loci (Figure 4a). Instead, our findings suggest that GCN5 indirectly regulates *PRX71* and *PRX33*, potentially through transcription factors such as *MYBS2* and *GATA21* (Figure 4b; Figure S3c). This model is consistent with our observation that H3K9ac is reduced at *PRX* promoters (Figure 4a) while *PRX71/PRX33* transcript abundance is elevated (Figure 1b), implying an indirect regulatory route rather than direct activation. However, genome-wide analysis has shown that loss of GCN5 leads to changes in H3K14ac enrichment along genes, with decreased H3K14ac at 5′ regions for GCN5 target genes but increased enrichment toward the 3′ ends of upregulated genes. Changes in H3K9ac closely follow those observed for H3K14ac along genes (Kim *et al*. 2020). Such changes in histone mark distribution could potentially influence H3K9ac at *PRX* loci and contribute to their transcriptional upregulation in *gcn5*, in parallel with the TF-mediated pathway. Evidence from rice supports a repressive role for *MYBS2*, as reduced *MYBS2* expression leads to elevated *αAmy* gene induction under abiotic stress conditions and improved stress tolerance (Chen *et al*. 2019). A similar model has been reported previously, in which GCN5 limits salicylic acid mediated responses by promoting the expression of genes that act as negative regulators of salicylic acid biosynthesis (Kim *et al*. 2020), suggesting that GCN5 may control biotic stress responses through induction of negative regulators. Together, these observations support a working model in which GCN5 maintains an acetylation-dependent transcriptional program that supports expression of *PRX* constraining TFs, thereby limiting stress-associated lignification and its growth trade-offs under salt stress (Figure 5). In the absence of GCN5, reduced H3K9ac at TF loci may lower TF abundance, relieving repression on *PRX71/PRX33* and promoting lignin deposition. This regulatory model is summarized in Figure 5. In wild type, *GCN5* promotes H3K9ac at loci encoding *PRX*, regulating transcription factors (*GATA21* and *MYBS2*), maintaining their expression. These TFs repress *PRX33*/*PRX71*, thereby limiting *PRX*-driven lignin deposition. In the *gcn5* mutant, reduced H3K9ac is associated with decreased TF expression, relieving repression of *PRX71*/*PRX33* and resulting in elevated PRX abundance and enhanced lignin deposition (Figure 5). This model provides an explanation for the apparent disconnect between reduced H3K9ac at *PRX* promoters (Figure 4a) and increased *PRX71*/*PRX33* transcript abundance (Figure 1b), and it is consistent with the ability of *PRX71/PRX33* overexpression to drive lignification under non-stress conditions (Figure 2) and with reduced salt-associated lignin deposition in *prx* mutants (Figure 3d).

**Figure 5.**
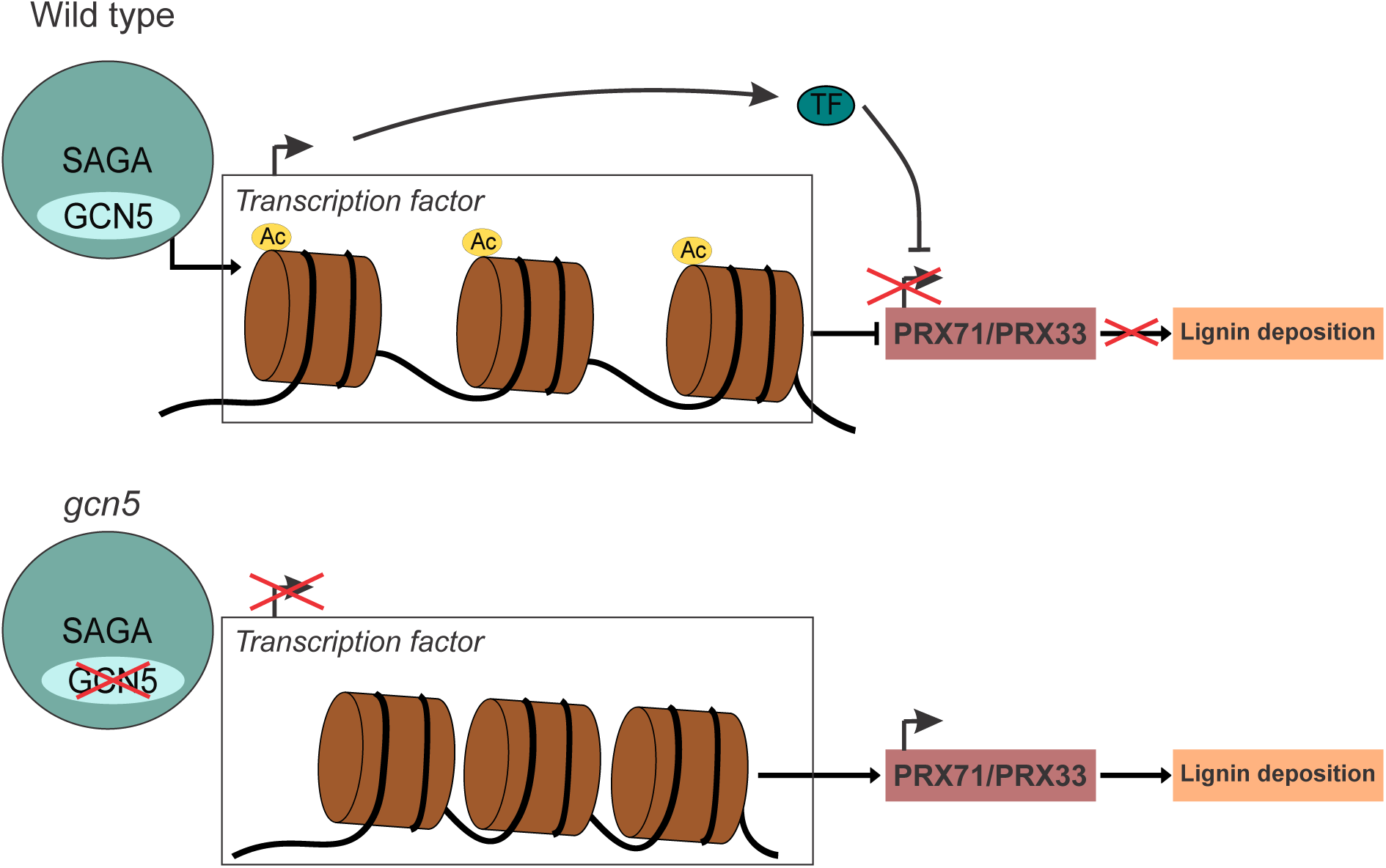
Proposed model for indirect regulation of *PRX71/PRX33* and lignin deposition by GCN5. In wild type (top), GCN5 promotes H3K9 acetylation (H3K9ac) at loci encoding *PRX* regulating transcription factors (TFs; e.g., *GATA21* and *MYBS2*), maintaining TF expression. These TFs repress *PRX71/PRX33*, thereby limiting *PRX* driven lignin deposition. In the *gcn5* mutant (bottom), reduced H3K9ac is associated with decreased TF expression, relieving repression of *PRX71/PRX33* and resulting in elevated *PRX* activity and enhanced lignin deposition.

Future experiments that directly test causality, for example, genetic epistasis between *gcn5* and *prx33/prx71*, quantitative ROS and lignin measurements beyond staining, and functional validation of *GATA21/MYBS2* at *PRX* promoters, will be important to distinguish primary chromatin-mediated effects from downstream stress responses and to refine the regulatory hierarchy. More broadly, these approaches will help clarify the contribution of *PRX71/PRX33* to the *gcn5* phenotype and their potential role in ROS and cell wall integrity signaling under salt stress.

## Acknowledgements

This work was supported by the Czech Science Foundation grant No. 22-17092S (PK and MW). We thank Dr. Fatemeh Aflaki for help with ChIP-qPCR, Adam Zeiner for valuable advice on physiological experiments, Media Kitchen (Tomas Kocabek) and the Growth Facility (Jan Kadlec), BC Core Facilities, Institute for Plant Molecular Biology, Biology Centre CAS.

## Competing interests

The authors declare no competing interests.

## Author contributions

MS, PK, IM and MW conceived the study and designed the research. MS and JM carried out the work. MS, JM, PK, IM and MW analyzed the data. MS wrote the first draft. All authors edited the final draft of the paper. PK and MW acquired funding for the research.

## Data availability

Data, plasmids and transgenic plant seeds generated for this study are available upon request from Michael Wrzaczek.

## Supplementary figure legends

**Supplementary Figure S1** Salt sensitivity and differentially expressed *PRX* genes in the *gcn5* mutant. (a) Survival rates of seedlings after salt treatment. Values are normalized to the corresponding genotype grown under control conditions. Data represent mean ± SE (n ≥ 3 biological replicates). Statistical analysis was performed using two-way ANOVA after confirming homogeneity of variance (Levene’s test, *p* > 0.05), followed by Tukey’s HSD test. Different letters indicate significant differences. (b) Primary root length (mm) measured at day 12 for seedlings grown on control or 150 mM NaCl-containing medium. Statistical analysis was performed using the Kruskal–Wallis test due to unequal variances (Levene’s test, p < 0.05), followed by the Wilcoxon post-hoc test. Different letters indicate significant differences. (c) Differentially expressed *PRX* genes in the *gcn5* mutant were identified from publicly available transcriptomic data. *PRX/PER* genes belong to class III peroxidases, whereas *GPX* genes encode glutathione peroxidases. (d) Expression levels of differentially expressed *PRX* genes at various time points during 150 mM NaCl treatment. Data represent mean ± SE of two biological replicates. Statistical significance was determined using two-way ANOVA followed by Tukey’s HSD test. Different letters indicate significant differences.

**Supplementary Figure S2** Salt stress responses in *gcn5*, complementation lines, and *prx* mutants. (a) Survival of plants grown on control and NaCl containing media. Scale bar = 1cm. (b) Primary root length (mm) measured at day 12 for seedlings grown on control or 150 mM NaCl containing medium. (c) Primary root length (mm) measured at day 12 for seedlings grown on control or 2.5 nM isoxaben containing medium. Data were analyzed using the Kruskal–Wallis test due to unequal variances (Levene’s test, *p* < 0.05), followed by the Wilcoxon post-hoc test. Different letters indicate significant differences.

**Supplementary Figure S3** Analysis of candidate transcription factors under control conditions. (a) Venn diagrams constructed using publicly available transcriptomic (RNA-seq) and ChIP-seq datasets. In (i), A represents genes downregulated in *gcn5* identified by RNA-seq (Wu *et al*. 2023), B represents genes showing reduced H3K9ac enrichment in *gcn5* identified by ChIP-seq (Wu *et al*. 2023), and C represents GCN5 target genes identified in wild-type plants by ChIP-seq (Kim *et al*. 2020). In (ii), D represents genes downregulated in *gcn5* identified by RNA-seq (Benhamed *et al*. 2008), E represents genes showing reduced H3K9ac enrichment in *gcn5* identified by ChIP-seq (Wu *et al*. 2023), and F represents GCN5 target genes identified in wild-type plants by ChIP-seq (Kim *et al*. 2020). In (iii), G represents genes downregulated in *gcn5* identified by RNA-seq (Wu *et al*. 2023), H represents genes showing reduced H3K9ac enrichment in *gcn5* identified by ChIP-seq (Wu *et al*. 2023), and I represents GCN5 target genes identified in wild-type plants by ChIP-seq (Wu *et al*. 2023). A downward arrow (↓) indicates lower enrichment. (b) Venn diagrams comparing transcription factor binding sites (TFBS) of the PRX71 (upper row) and PRX33 (lower row) promoters with overlapping gene sets from (a) i–iii. Letter combinations (e.g., B ∩ C, A ∩ B, E ∩ F) indicate intersections between the gene sets identified in (a) i–iii. Only combinations with at least one common gene are shown. Combinations with zero overlapping genes were observed but not shown above. TFs obtained from the gene set intersections were compiled. After removing duplicates, 16 TFs were selected for further analysis. These TFs include those shared between the PRX71 and PRX33 promoters, as well as those specific to either promoter. (c) Expression levels of differentially expressed transcription factor genes under control conditions measured by qRT–PCR. Data represent mean ± SD of three biological replicates. Statistical significance was determined using a two-tailed Student’s t-test.

**Supplementary Figure S4** Confirmation of T-DNA mutant (a) Genomic DNA from Col-0 and mutant plants was used as a template for PCR to confirm the presence of T-DNA insertion lines. (b) RT-PCR analysis showed the absence of transcript in the mutants. *GCN5* specific primers were used in lanes 1 and 2, *PRX71-1* specific primers in lanes 6 and 7, *PRX71-2* specific primers in lanes 11 and 12, and *PRX33* primers in lanes 16 and 17. *PP2AA3* was used as an internal reference for all samples (lanes 3, 4, 8, 9, 13, 14, 18, and 19).

**Supplementary Table S1.** Primer sequences used for all experimental analyses.

**Supplementary Table S2.** Potential transcription factor binding sites in the promoters of *PRX71* and *PRX33*.

**Supplementary Table S3.** Differentially expressed genes (DEGs) and ChIP peak changes in selected genes.

## Notes

### Competing Interest Statement

The authors have declared no competing interest.

### Summary of Updates

We have updated the analyses of the transcription factors putatively regulating PRX expression.

