## Supplementary Figures for "GCN5 restricts PRX-dependent lignin deposition during salt stress in Arabidopsis"

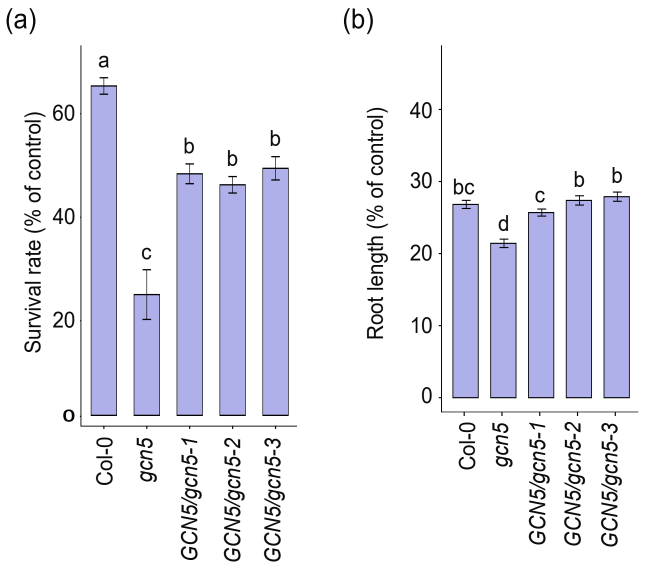


(c)

| Gene ID | Gene Name | Regulation |
|---|---|---|
| AT1G05250 | PRX2 | Up |
| AT1G14550 | PER5 | Up |
| AT1G30870 | PER7 | Up |
| AT1G68850 | PER11 | Up |
| AT2G37130 | PER21 | Up |
| AT2G39040 | PER24 | Up |
| AT3G49110 | PRX33 | Up |
| AT3G49120 | PRX34 | Up |
| AT3G49960 | PER35 | Up |
| AT4G16270 | PER40 | Up |
| AT4G30170 | PER45 | Up |
| AT4G33870 | PER48 | Up |
| AT4G36430 | PER49 | Up |
| AT5G05340 | PRX52 | Up |
| AT5G24070 | PER61 | Up |
| AT5G39580 | PER62 | Up |
| AT5G64100 | PER69 | Up |
| AT5G64110 | PER70 | Up |
| AT5G64120 | PRX71 | Up |
| AT2G48150 | GPX4 | Up |
| AT2G41480 | PRX25 | Down |
| AT2G25080 | GPX1 | Down |
| AT4G31870 | GPX7 | Down |

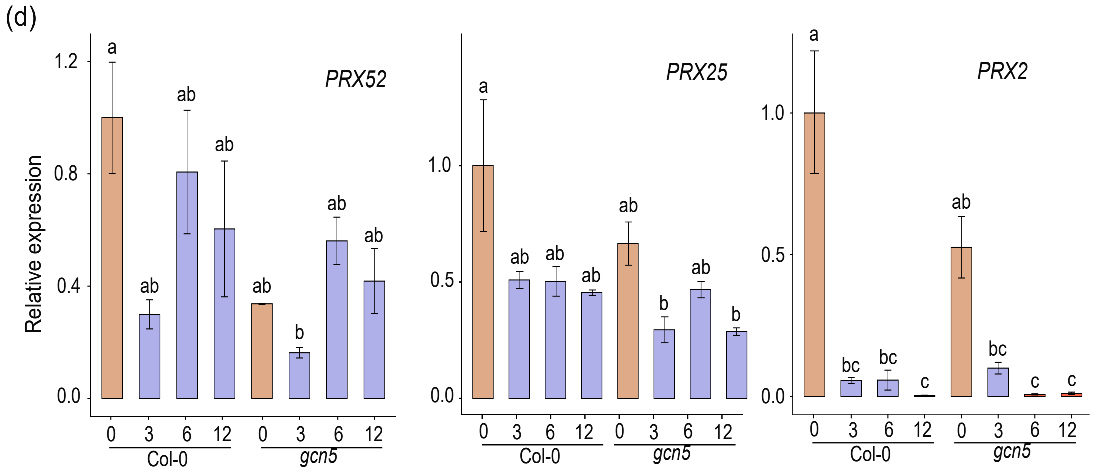


**Figure S1** Salt sensitivity and differentially expressed *PRX* genes in the *gcn5* mutant. (a) Survival rates of seedlings after salt treatment. Values are normalized to the corresponding genotype grown under control conditions. Data represent mean ± SE (n ≥ 3 biological replicates). Statistical analysis was performed using two-way ANOVA after confirming homogeneity of variance (Levene's test, $p > 0.05$), followed by Tukey's HSD test. Different letters indicate significant differences. (b) Primary root length (mm) measured at day 12 for seedlings grown on control or 150 mM NaCl-containing medium. Statistical analysis was performed using the Kruskal–Wallis test due to unequal variances (Levene's test, $p < 0.05$), followed by the Wilcoxon post-hoc test. Different letters indicate significant differences. (c) Differentially expressed *PRX* genes in the *gcn5* mutant were identified from publicly available transcriptomic data. *PRX/PER* genes belong to class III peroxidases, whereas *GPX* genes encode glutathione peroxidases. (d) Expression levels of differentially expressed *PRX* genes at various time points during 150 mM NaCl treatment. Data represent mean ± SE of two biological replicates. Statistical significance was determined using two-way ANOVA followed by Tukey's HSD test. Different letters indicate significant differences.

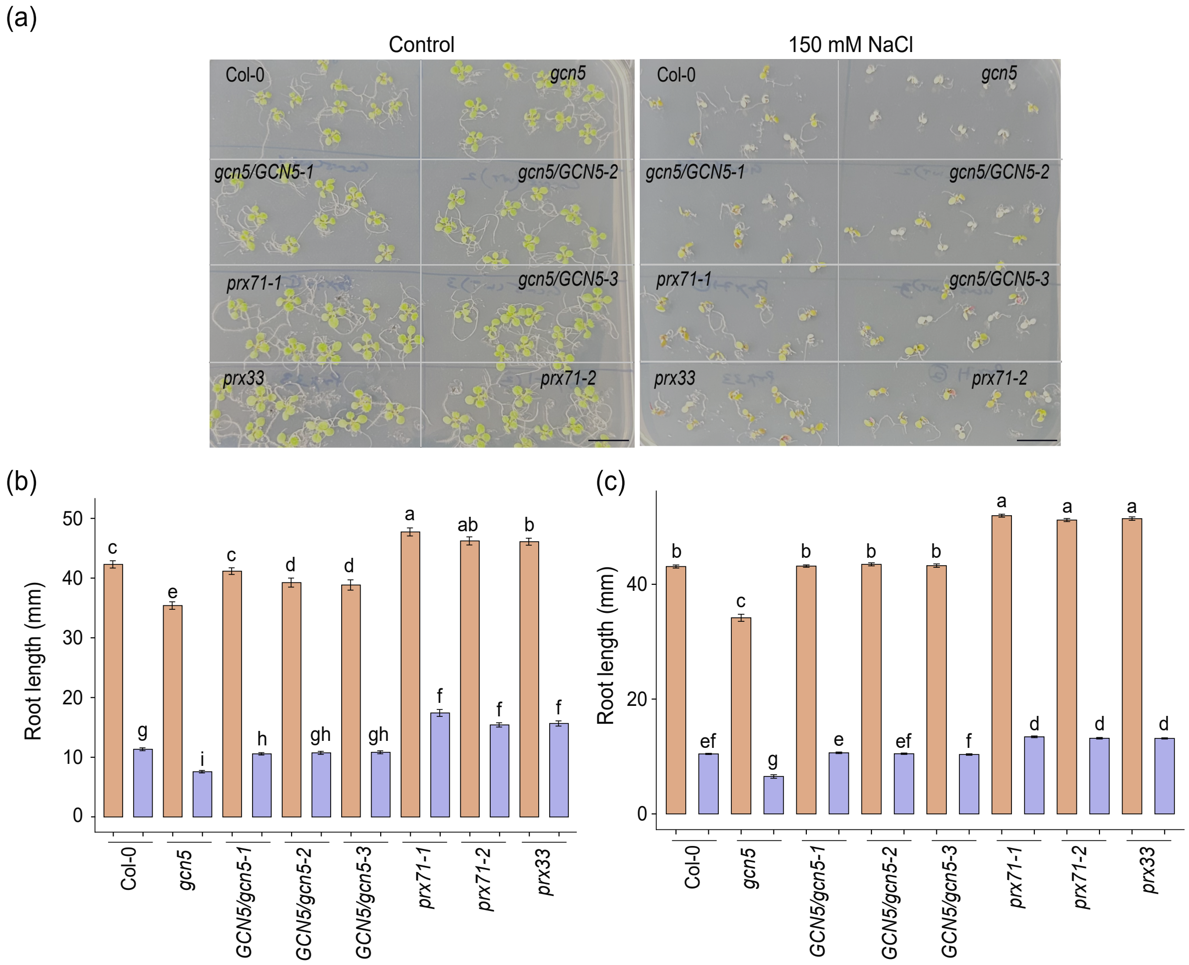


**Figure S2** Salt stress responses in *gcn5*, complementation lines, and *prx* mutants. (a) Survival of plants grown on control and NaCl containing media. Scale bar = 1cm. (b) Primary root length (mm) measured at day 12 for seedlings grown on control or 150 mM NaCl containing medium.(c) Primary root length (mm) measured at day 12 for seedlings grown on control or 2.5 nM isoxaben containing medium. Data were analyzed using the Kruskal–Wallis test due to unequal variances (Levene's test, $p < 0.05$), followed by the Wilcoxon post-hoc test. Different letters indicate significant differences.

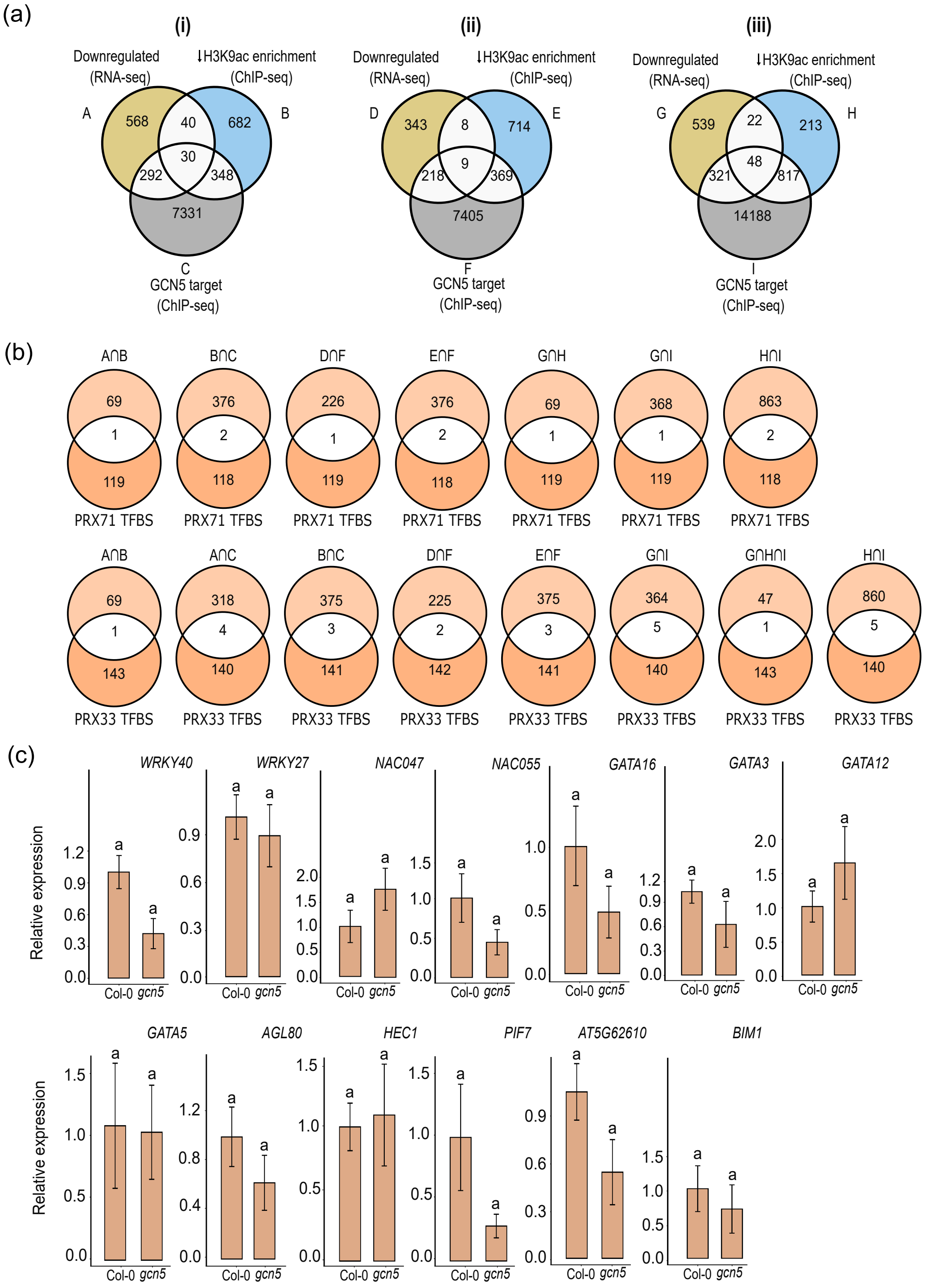


**Figure S3** Analysis of candidate transcription factors under control conditions. (a) Venn diagrams constructed using publicly available transcriptomic (RNA-seq) and ChIP-seq datasets. In (i), A represents genes downregulated in *gcn5* identified by RNA-seq (Wu et al. 2023), B represents genes showing reduced H3K9ac enrichment in *gcn5* identified by ChIP-seq (Wu et al. 2023), and C represents GCN5 target genes identified in wild-type plants by ChIP-seq (Kim et al. 2020). In (ii), D represents genes downregulated in *gcn5* identified by RNA-seq (Benhamed et al. 2008), E represents genes showing reduced H3K9ac enrichment in *gcn5* identified by ChIP-seq (Wu et al. 2023), and F represents GCN5 target genes identified in wild-type plants by ChIP-seq (Kim et al. 2020). In (iii), G represents genes downregulated in *gcn5* identified by RNA-seq (Wu et al. 2023), H represents genes showing reduced H3K9ac enrichment in *gcn5* identified by ChIP-seq (Wu et al. 2023), and I represents GCN5 target genes identified in wild-type plants by ChIP-seq (Wu et al. 2023). A downward arrow (↓) indicates lower enrichment. (b) Venn diagrams comparing transcription factor binding sites (TFBS) of the PRX71 (upper row) and PRX33 (lower row) promoters with overlapping gene sets from (a) i–iii. Letter combinations (e.g., B ∩ C, A ∩ B, E ∩ F) indicate intersections between the gene sets identified in (a) i–iii. Only combinations with at least one common gene are shown. Combinations with zero overlapping genes were observed but not shown above. TFs obtained from the gene set intersections were compiled. After removing duplicates, 16 TFs were selected for further analysis. These TFs include those shared between the *PRX71* and *PRX33* promoters, as well as those specific to either promoter.
(c) Expression levels of differentially expressed transcription factor genes under control conditions measured by qRT–PCR. Data represent mean ± SD of three biological replicates. Statistical significance was determined using a two-tailed Student's t-test.

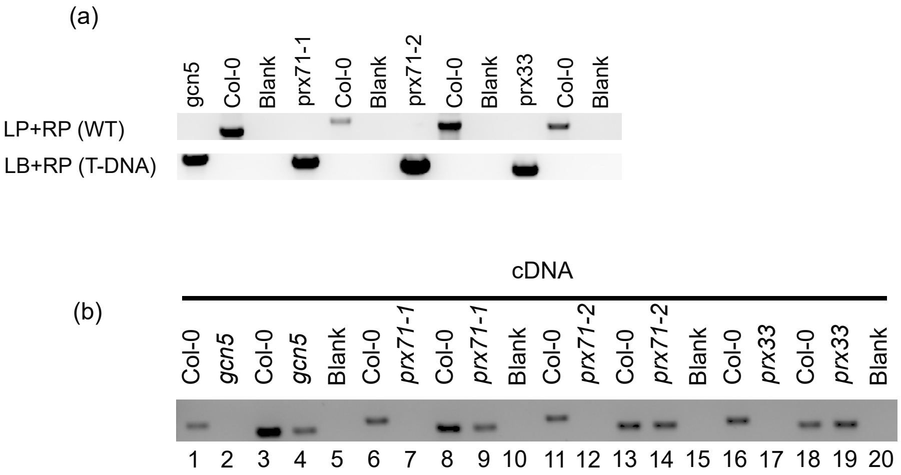


**Figure S4** Confirmation of T-DNA mutant (a) Genomic DNA from Col-0 and mutant plants was used as a template for PCR to confirm the presence of T-DNA insertion lines. (b) RT-PCR analysis showed the absence of transcript in the mutants. *GCN5* specific primers were used in lanes 1 and 2, *PRX71-1* specific primers in lanes 6 and 7, *PRX71-2* specific primers in lanes 11 and 12, and *PRX33* primers in lanes 16 and 17. *PP2AA3* was used as an internal reference for all samples (lanes 3, 4, 8, 9, 13, 14, 18, and 19).
